# Forest habitat and forest dominated landscapes are associated with bumblebee species with visual traits related to light sensitivity

**DOI:** 10.1101/2024.12.14.628511

**Authors:** Océane Bartholomée, Pierre Tichit, Jens Åström, Henrik G. Smith, Sandra Åström, Markus A. K. Sydenham, Emily Baird

## Abstract

While functional traits like body size have been extensively linked to species distributions, the influence of sensory traits on species’ responses to environmental changes remains underexplored. Particularly, the relationship between light sensitivity and niche segregation across different distributional extents – local habitat conditions and across entire landscapes – remains unclear. In this study, we examined bumblebee communities monitored across Norway on grassland and forest habitats within landscapes varying in forest cover within 1 km radii. We investigated whether the eye parameter – a visual trait measuring the trade-off between light sensitivity (high values) and visual resolution (low values) – was associated with local habitat types and the forest cover at the landscape scale. Additionally, we combined bumblebee-plant interactions with a plant trait, to determine if bumblebee light sensitivity correlated with the shade tolerance of the plants they foraged on. Our findings showed that bumblebee species with high eye parameters were more common and abundant in forest habitats and areas with greater forest cover, while species with low eye parameters showed the opposite trend. This pattern was also reflected at the community level, as indicated by the community-weighted mean of the eye parameter which increased with forest cover and was higher in forest habitats. Furthermore, bumblebees with higher eye parameters tended to forage on plants with greater shade tolerance.

These results suggest that visual adaptations for light sensitivity contribute to shaping bumblebee species distributions across different scales. Overall, our study underscores the importance of pollinator vision in understanding species niches, in relation to habitat use and foraging behaviour. By relating pollinator visual abilities to plant niches for the first time, this study provides an important basis for future modelling of plant-pollinator interactions and targeted conservation measures for both plants and pollinators in forested landscapes.

## Introduction

Pollinator declines threaten ecosystem functioning because the reproductive success of the vast majority of flowering plants benefits from animal pollination (Ollerton et al. 2011). A major driver of recent pollinator declines including bees, is human-driven changes of habitat availability and quality, resulting in loss of foraging resources and nesting sites (Potts et al. 2010; IPBES 2016). To increase the capacity to predict pollinator responses to environmental changes, it is necessary to determine how habitat characteristics – e.g temperature and light conditions – influence pollinator species distributions (Williams et al. 2010). Species sharing ecological traits that affect their fitness can be expected to respond similarly (McGill et al. 2006), so that species-sorting along environmental gradients operates through ‘trait-based environmental filtering’ (Vellend & Agrawal 2010).

Trait-based approaches are an increasingly used analytical pathway that uses functional traits to describe and predict species responses to global changes (Green et al. 2022) in an attempt to identify mechanistic links between community dynamics and environmental conditions (McGill et al. 2006). The traits of insect pollinators, such as bees, can be broadly classified into: autecological traits describing the life histories of species; locomotive ability traits; and climatic tolerance traits (Martinet et al. 2021). A fourth category are sensory traits that describe how pollinators are able to respond to the sensory cues necessary for finding plants (Spaethe et al. 2001) and interacting with their environment (Bartholomée et al. 2023). For insect pollinators, body size (Green et al. 2022) is widely used as a functional trait, making mechanistic interpretation hard as it is tightly correlated to several functions. Body size of bees, when used as a proxy for thermoregulatory abilities, has been related to altitudinal gradients (Peters et al. 2016; Osorio-Canadas et al. 2022) and, when used as a proxy for mobility, mediates responses to habitat fragmentation (Carrié et al. 2017; Gérard et al. 2021). Therefore, it is difficult to relate changes in the mean body size of species within a community to a specific ecological mechanism. Analyses of trait-environment relationships have taught us how environmental conditions filter for traits related to mobility, as well as resource use (Hoiss et al. 2012; Sydenham et al. 2015). However, we lack a clear understanding of how insect pollinator species differ in their ability to navigate and forage efficiently under changes in light conditions, as being active under a forest canopy in dim light can be visually challenging (Arnold & Chittka 2012).

Sensory traits dictate how animals experience and interact with their environment (Dominoni et al. 2020; Elmer et al. 2021). Vision is essential, since highly mobile animals like flying insects need to orient themselves in habitats, to find flowers (Spaethe et al. 2001) and – for central place foragers such as bees – to commute between their nests and food resources (Warrant et al. 2004). Recent studies have related eye size to light microhabitat use for damselflies (Scales & Butler 2016) and butterflies (Wainwright et al. 2023), but not in bumblebees (Bartholomée et al. 2023). In contrast, the visual traits of diurnal bumblebee species more subtly influence their flight activity at different light intensities (Kaputjanski et al. 2007). Variations in a visual trait known as the eye parameter – a measure of how an insect compound eye balance light sensitivity, which is important in dim light conditions, and visual resolution, which is possible under bright conditions (Jander & Jander 2002) (Figure 1a) – explain how different bumblebee species use light microhabitats in hemi-boreal forest understory at the scale of a light patch on the understory floor (*i.e.*, the scale at which these pollinators forage) (Bartholomée et al. 2023). How sensory traits are associated with variation in pollinator communities at larger scales and across wider geographical ranges remains unclear. A recent study focusing on grasslands across Sweden found that bumblebee communities in landscapes with a higher forest cover had higher eye parameter, suggesting enhanced visual abilities in the challenging light conditions of a forest (Tichit et al. under review). Nevertheless, a more in-depth investigation of potential differences between scales, that is to say if that pattern is consistent across local habitats – *i.e.*, by including communities sampled both in grasslands and forests – remains to be investigated. Such analyses can indicate if bumblebee species are able to navigate in contrasted habitats but do not provide information if these species are able to forage in these contrasted light habitats, as vision is involved both in microhabitat use (Bartholomée et al. 2023) and flower detection (Spaethe et al. 2001). This can be investigated by testing if the niche of plant communities foraged by bumblebees are consistent with their visual abilities – strengthening the visually mediated community assembly.

**Figure 1:**
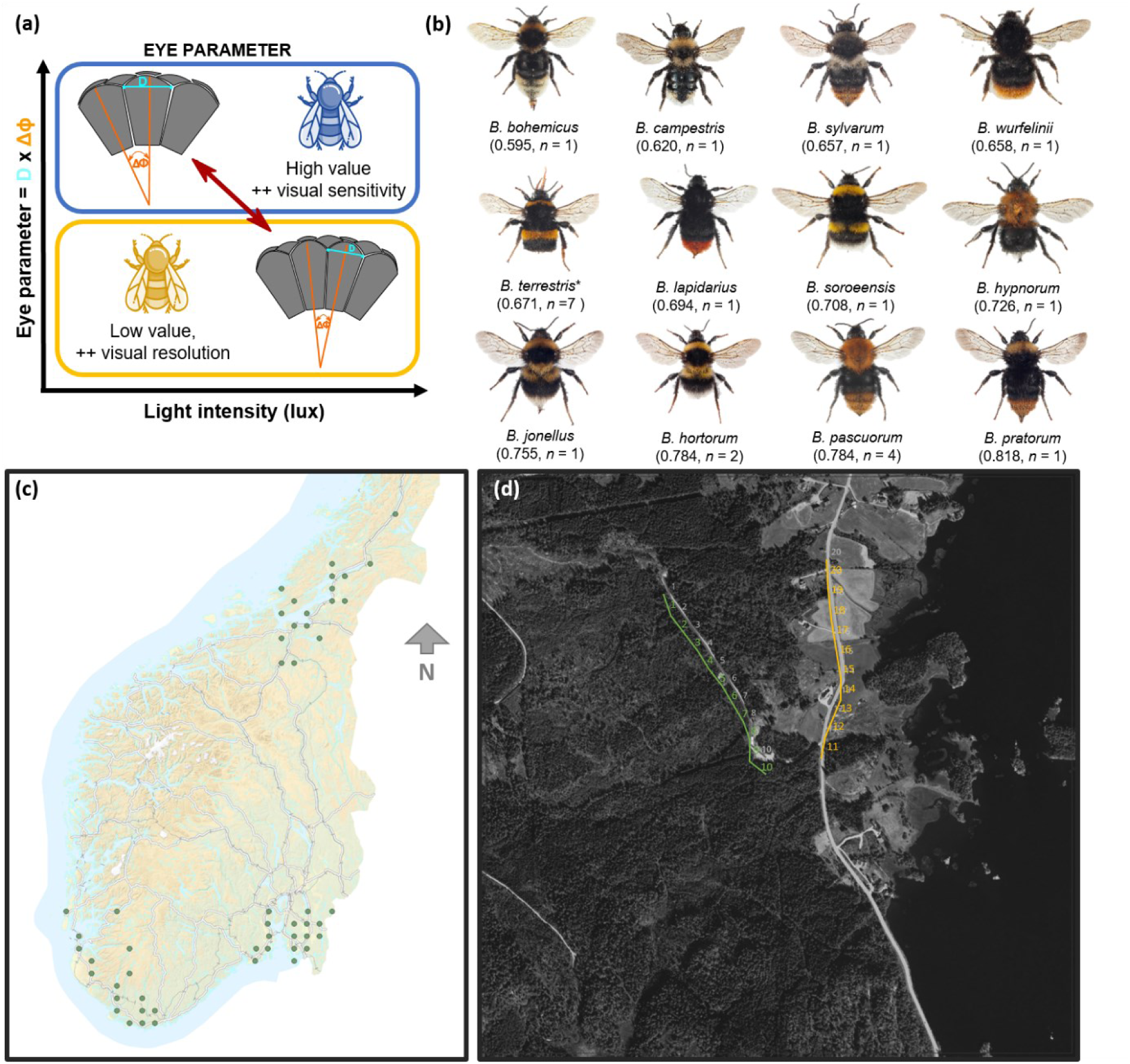
(a) The eye parameter reflects the trade-off between spatial resolution and light sensitivity in insect compound eyes. To cope with reduced light intensity (top left), insects living in dim habitats typically have higher facet diameters (D, μm) and higher angle between each unit (Δφ, rad) than their bright light counterparts (bottom right). (Adapted from Bartholomée et al. (2023)). (b) 11 bumblebee species from the Norwegian national inventory included in the analysis of question 1 (pictures not on scale). Species are sorted by increasing values of eye parameter. Each picture is annotated with the species name (*B.* is for *Bombus*) (CC BY 3.0, © Arnstein Staverløkk). In between bracket: species-specific eye parameter value (μm.rad) and sample size of the measurements (*n*). \**B. terrestris* includes here both *B. lucorum* and *B. terrestris*, as both species are highly difficult to distinguish in the field (Carolan et al. 2012). (c) Map of the monitored sites in Norway. (d) Close-up of survey square 803 in the South-Western region, with numbered open forest transects in green and grassland transects in yellow.

Here we test if forests are associated with bumblebee species with higher light sensitivity, which we hypothesise, increases their relative fitness in shaded microhabitats. We do this at the local scale by contrasting forest vs. grassland habitats, and at the landscape scale by comparing landscapes with varying overall percentage cover of forest. We used data on bumblebee species distributions from long-term structured surveys performed in both grasslands and open forests (Åström & Åström 2023). The eye parameter trait was measured on a low number of bumblebee individuals sampled outside the study monitoring range (Tichit et al. under review), due to the complexity and the time required obtain this calculation (Taylor et al. 2019). To strengthen our hypothesis, we tested if bumblebees with a better dim light vision have a stronger affinity for shade-tolerant plants than species with a higher visual resolution, which could indicate that these bumblebee species are able to both navigate and forage in dim light environments. In order to understand the interplay between bumblebee vision and the light niche they forage on, we focused on plant-bumblebee interactions extracted from an open-source database (Database of Pollinator Interactions) (Balfour et al. 2022) combined with a trait database gathered for Swedish plants species (Tyler et al. 2021).

## Material and Methods

### Bumblebee monitoring data

Since 2009, the Norwegian Institute for Nature Research (NINA) has managed a (soon to be) country-wide bumblebee monitoring program in both open lowland – defined as open areas below the treeline – called here grasslands for simplicity, and forests, with data made available on GBIF (Åström & Åström 2023). We focused on the monitoring done over nine years of sampling from 2013 until 2021, *i.e.* when the geographic span of the project was extended to three geographical regions (58.06°-64.93°N, 5.22°- 13.18°W, Figure 1c). Monitoring was performed in 52 sites at low altitude (mean [min – max]: for open low land sites: 110 [0 – 487] m.a.s.l ; for forest sites: 195 [3 – 722] m.a.s.l.), each corresponding to a 1.5 x 1.5 km square located at least at 18 km from the other sites. In each square, 1 km of transect (20 x 50 m transects) is walked three times per year. Within each geographical region, the total number of transects are divided into roughly equal parts between open forest and grassland habitats (Figure 1d). The transects are walked between 10:00 and 17:00 in suitable weather conditions for insect activity (> 15°C, < 60% cloud cover, weak wind). Bumblebees are counted and identified visually within 2.5 m on each side of the transect and 5 m ahead. We focused our analysis on the 12 most abundant species (*i.e.,* species with ≥ 30 individuals observed in the nine years), 10 non-cuckoo species and two cuckoo species (*B. bohemicus* and *B. campestris*) (Figure 1b), resulting in a dataset of 47 sites. Although cuckoo and non-cuckoo species differ in their life history traits, we include them together because they must both have sensory traits that allow them to be active in the habitats they are observed (Table S1). The species reported as *Bombus* sensus stricto was interpreted as the *B. terrestris* complex, which can include *B. terrestris, B. lucorum, B. magnus* and *B. cryptarum,* which are difficult to separate in the field (Carolan et al. 2012).

### Landscape composition

We used the Copernicus CLC+ Backbone (10x10 m, from 2018), which include 11 basic land cover classes (European Union’s Copernicus Land Monitoring Service information 2022) to estimate the forest cover (in %) in a 1 km radius buffer around the centroid of each transect – as most bee species have a flight range below 1 km (Kendall et al. 2022). Forest cover was calculated as the sum of the percentage cover (%) of both coniferous (land cover class 2 “Woody – needle leaved trees”) and broadleaf (land cover class 3 “Woody – broadleaved deciduous trees”) forests using the exact_extract function in the *exactextract* package (Baston 2022). The GIS analyses were performed in R v.4.3.0 (R Core Team 2023) with the packages *raster* (Hijmans 2023), *sf* (Pebesma & Bivand 2023) and *exactextractr* (Baston 2022).

### Visual trait measurements

To study how the visual abilities of bumblebee species might explain their relationships with their habitat, their landscape and the plants they forage on, we used the eye parameter, a measure reflecting the trade-off between light sensitivity and visual resolution (Jander & Jander 2002) (Figure 1a). Whereas eye and body sizes are allometrically related (Jander & Jander 2002; Howland et al. 2004), the eye parameter is only weakly related to body size, with correlations driven by within species differences (Tichit et al. under review). For example, in *B. terrestris,* a species with a high intra-specific body size variation, eye parameter increased with eye volume at an exponent of 0.6 (Fig. 3 in Taylor et al. 2019). In this study, we use values from species averages for individuals obtained from Tichit et al. (under review). In that study, the eye parameter was estimated from bumblebees sampled outside the monitoring range of this work, with average measurements taken across one eye of each individual. It was calculated by multiplying the facet diameter (in μm) by the average angle between the centre of two adjacent facets (in rad). For investigating how the eye parameter interacts with habitat and landscape forest cover, we included values from workers for the non-cuckoo species and queens for the cuckoo species, as this was the most common caste observed during the monitoring (Table S1). Due to the logistic difficulties in measuring the eye parameter, we only had one or a few measures on workers for each species (range: 1-7, mean: 1.7; details in Table S1). However, for 13 species where measures of at least three individuals were available, the proportion of variance explained by species identity was 63 % (linear model of eye parameter explained by species) (Tichit et al., submitted). Moreover, for the species/caste included in this study, eye parameter was not related to body size (linear mixed model: eye parameter ∼ ITD + (1|species)). To relate bumblebee visual abilities to the light niche of the plants they forage on, we used the eye parameters of workers for bumblebee species (11 species, 10 from the previous analyses and one less abundant species) – and the queen trait for the cuckoo species (three species, one from the previous analyses and two less abundance species). In total, we included 15 species (Table S1).

### Body size – Intertegular distance (ITD)

To control for potential effect of body size on species responses across scales, we compiled and averaged ITD measures from several literature sources for the workers of non-cuckoo species and the queens of the cuckoo species (Supporting information, Table S2) (Peat et al. 2005; del Castillo & Fairbairn 2012; Streinzer & Spaethe 2014; Kendall et al. 2019; Massa et al. 2024; Tichit et al. under review). For the *B. terrestris* complex, we averaged the values available for both *B. terrestris* and *B. lucorum* (Table S2).

### Plant-bumblebee interactions and plant light niche

To obtain data on bumblebee-plant interactions, we used DoPi (Database of Pollinator Interactions) (Balfour et al. 2022), which focuses on pollinators in the United Kingdom – including 14 out of the 20 bumblebee species encountered in the Norwegian bumblebee monitoring data – to which we added *B. subterraneus*, as its eye parameter was available. We downloaded the data for each species. For the *B. terrestris* complex, the selection was done through the option “*Bombus lucorum/terrestris*”. We first included all the plants for which any interaction has been reported in the DoPI database. To refine our search to Scandinavian plant taxa, we combined this data with a plant trait database compiled for Swedish plant species (Tyler et al. 2021). We focused on the trait “light index”, which is a measure of the “optimal light/shade conditions of the species” and is a semi-quantitative measure going from 1 (deep shade) to 7 (always full sun), based on expert knowledge and flora atlases.

The raw plant data of the interactions reported in DoPi included 853 plant taxa – *i.e.* species and genus when a species was reported as “spp.”. We included all unique bumblebee species-plant taxon interactions from the database. For plant taxa reported at the genus level, we took the average light index trait values across species of that genus. From these 853 taxa, 596 were initially found in the Swedish database. A manual search for taxonomic synonyms of the 257 missing taxa allowed to add an extra 19 taxa, resulting in the inclusion of 615 plant taxa in the analyses. Plant-bumblebee interactions related to the plant taxa absent of the Swedish plant trait database were removed. The numbers of unique bumblebee species-plant taxa associations are detailed in the Table S1.

### Statistical analysis

All statistical analyses were performed with the statistical software R (R Core Team 2023). Generalised linear mixed models (GLMMs) were fitted with the *glmmTMB* package (Brooks et al. 2017). Diagnostics of their residuals were checked with the *DHARMa* package *(Hartig 2022)*. The post-hoc contrast analyses were performed with *emmeans* (Lenth 2023). Plots to visualise results were made with *ggplot2* (Wickham 2016) and *sjPlot* (Lüdecke 2024).

#### Are forest habitat and landscape forest cover that reduce light availability associated with pollinator species with high light sensitivity?

To test if visual abilities influence how bumblebees interact with local habitat and landscape characteristics, we pooled the community data per transect across the nine years of sampling to exclude inter-annual variation in community composition and to overcome the zero inflation of the yearly data – resulting in 1081 transects. We transformed the data into ‘long format’, to get one species observation per transect per row, resulting in 12972 observations (12 species x 1081 transects). In these analyses, we refer to the percent of forest at the landscape scale as ‘forest cover’ and the local habitat type of the individual transects as ‘forest habitat’ and ‘grassland habitat’. We built two GLMMs evaluating the effects of both local and landscape characteristics simultaneously, described in Table 1. The responses variables for Models 1 and 2 are respectively the probability of presence and the abundance of bumblebee species. For Model 2, a negative binomial distribution with a quadratic parametrisation was chosen to overcome over-dispersion. In Models 1 and 2, we accounted for variation in sampling effort by including logged sampling effort respectively as a covariate – as offsets are not implemented for binomial models – and as an offset. In both models, we included the sampling site as random effect to account for potential spatial autocorrelation. We controlled for geographical variation by integrating latitude, longitude, and their interaction as fixed effects. We accounted for the potential role of species body size on their response to forest cover and their habitat by including ITD in interaction with both environmental variables.

**Table 1:** Description of the four statistical models: two generalised linear mixed models (Models 1 and 2) and two linear mixed models (Models 3 and 4). The explanatory variables are: (i) species occurrence (binomial, 1/0), (ii) species abundance (integer), (iii) community weighted mean of the eye parameter (CWM, μm.rad), and (iv) the light index of foraged plant taxa (discrete, 1 to 7). The explanatory variables are: (i) eye parameter of each species (μm.rad), (ii) inter-tegular distance of each species (ITD, mm), (iii) habitat type (categorical, forest/grassland), (iv) forest cover in a 1 km radius (continuous, %), and (v) sampling effort is the number of visits (integer). In Models 1-3, we fitted the identity of the 47 sites of sampling as random effect, to control for potential spatial autocorrelation between transects in Models 1-3.

| Model (distribution) | Response variable | Formula | Random effect | Data sources |
| --- | --- | --- | --- | --- |
| Model 1<br>(binomial) | Species occurrence | eye parameter*(habitat type + forest cover) + ITD*(habitat type + forest cover) + longitude*latitude + log(sampling effort) | + (1\|Site ID) | Norwegian bumblebee monitoring data<br>Eye parameter from (Tichit et al. under review) |
| Model 2<br>(negative binomial with quadratic parametrisation) | Species abundance | eye parameter*(habitat type + forest cover) + ITD*(habitat type + forest cover) + longitude*latitude + offset(log(sampling effort)) | + (1\|Site ID) | Norwegian bumblebee monitoring data<br>Eye parameter from (Tichit et al. under review) |
| Model 3<br>(gaussian) | CWM eye parameter | habitat type + forest cover + longitude*latitude | + (1\|Site ID) | Norwegian bumblebee monitoring data<br>Eye parameter from (Tichit et al. under review) |
| Model 4<br>(gaussian) | Plant light index | eye parameter |  | Plant-bumblebee interaction from United Kingdom (Balfour et al. 2022)<br>Eye parameter from (Tichit et al. under review)<br>Plant light index from Swedish data base (Tyler et al. 2021) |

For Models 1 and 2, we performed post-hoc contrast analyses on the significant interaction terms with the emtrends function of the *emmeans* package (Lenth 2023). For the interaction between the eye parameter (and ITD) and forest cover, we focused on the contrast between the slopes of three values of eye parameter: high (average + SD), intermediate (average), low (average – SD). For the interaction between the eye parameter and habitat, we compared the slopes estimates of forest cover for the two habitat types.

Furthermore, to test if the observed patterns were consistent at the community level, we calculated the community-weighted mean (CWM) of the eye parameter for the community averaged across the years at the transect level – *i.e.,* the averaged trait value weighed by the abundance of each bumblebee species:

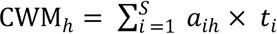

with *aih* the abundance of species *i* across of the 10 years of sampling *h* and *ti* the eye parameter value of the bumblebee species *i*. For that we used the function functcomp of the *FD* package (Laliberté & Legendre 2010; Laliberté et al. 2014). We built Model 3 described in Table 1.

#### Plants and bumblebees – Do bumblebees with a high light sensitivity predominantly visit shade-tolerant species?

To answer this question, we tested if the shade tolerance of the plant taxa visited by each bumblebee species is related to the light sensitivity of the bumblebees, as measured by the eye parameter. That is to say, we investigated if bumblebee species with a high light sensitivity (*i.e.*, having better dim light visual abilities) predominantly forage on plants with a higher shade tolerance (*i.e.*, growing in dimmer light conditions). To do this, we built the linear Model 4 with the light index of each of the plant taxa visited by individual bumblebee species as response variable and the bumblebee species eye parameter as explanatory variable. As each bumblebee species-plant taxa was unique, we did not include a random effect.

## Results

### Forest habitat and landscape forest cover that reduce light availability are associated with pollinator species with high light sensitivity

The occurrence and abundance of bumblebee species was higher in grassland than in forest habitats (occurrence: *Z* = 4.21, *P* < 0.001; abundance: Z = 3.73, *P* < 0.001), and decreased with increasing forest cover (occurrence: *Z* = -5.59, *P* < 0.001; abundance: Z = -8.75, *P* < 0.001) (Supporting information, Figure S1). Species with a high eye parameter (indicating a greater light sensitivity) were consistently more common than species with a lower eye parameter (occurrence: *Z* = 6.96, *P* < 0.001) while smaller species where more abundant than larger species (Figure S1). Our hypothesis was evaluated by post-hoc contrasts, which were consistent for the occurrence and abundance. The slopes of both habitats – forests and grassland habitats – were different for both occurrence and abundance (occurrence: *Z* = 4.78, *P* < 0.001; abundance: Z = 4.14, *P* < 0.001), with both occurrence and abundance increasing faster with eye parameter in forest habitats (Figure 2a, 2b, Supporting information, Table S3). The analyses showed that both occurrence and abundance of bumblebee species with a high eye parameter increased along a forest cover gradient (occurrence: *Z* = - 4.74, *P* < 0.001; abundance: Z = - 8.35, *P* < 0.001) (Figure 2c, 2d, Table S3). In contrast, the occurrence and abundance of bumblebee species with a low eye parameter decreased along the same forest cover gradient, while occurrence of species with intermediate eye parameter did not change with forest cover (95 % CI overlapping with 0) (Table S3). The occurrence and abundance of large bumblebee species increased with forest cover while the occurrence and abundance of small species decreased along the same gradient (occurrence: *Z* = 5.94, *P* < 0.001; abundance: Z = 6.75, *P* < 0.001) (Figure 2E, 2F, Table S3). The occurrence and, marginally, the abundance, of large species increased faster in forest than in grassland habitats (occurrence: *Z* = 1.97, *P* = 0.049; abundance: *Z* = 1.92, *P* = 0.055) (Table S3).

**Figure 2:**
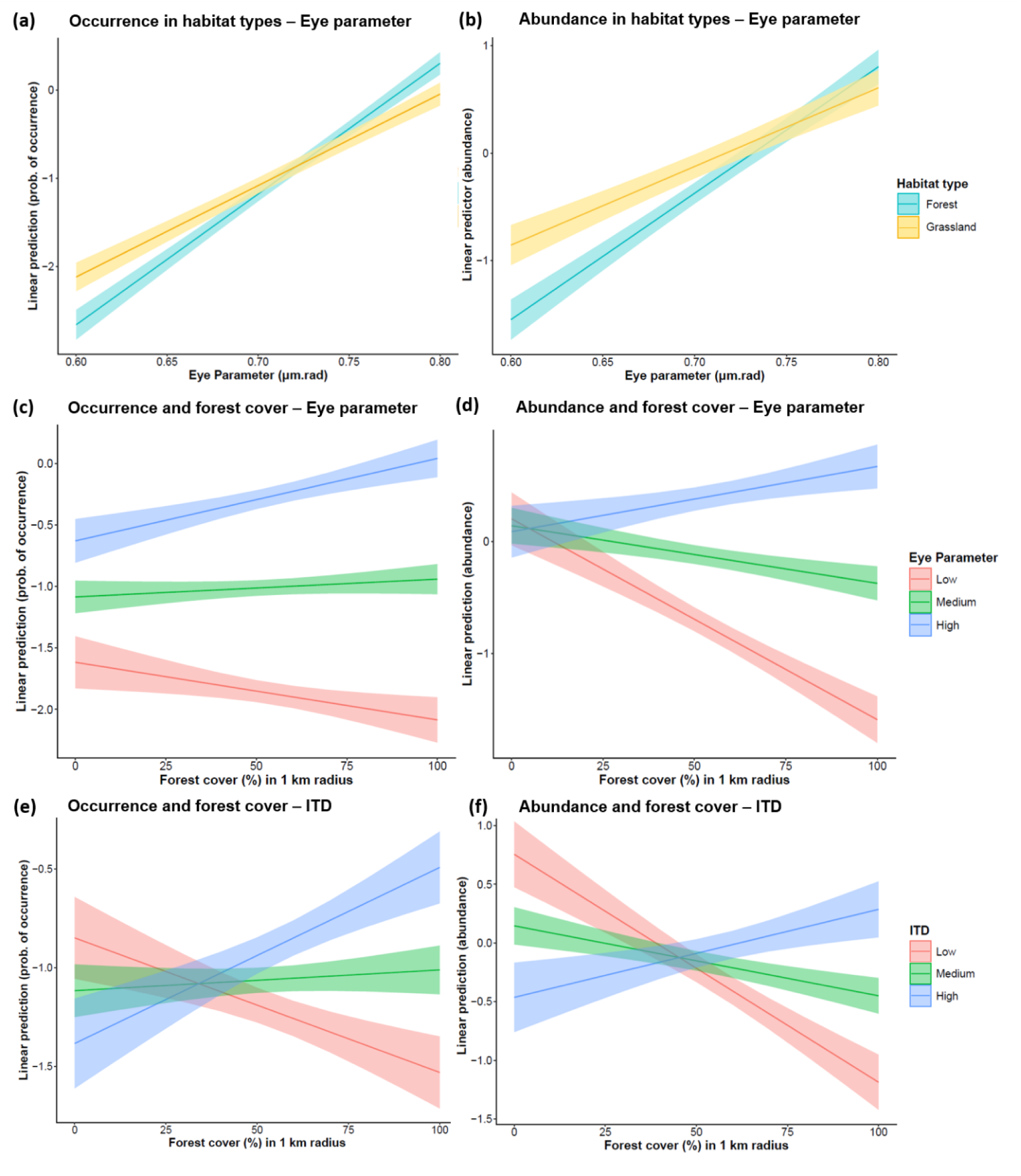
Post-hoc contrast analyses of the modelling of bumblebee communities (Models 1 and 2). The interaction between the eye parameter of bumblebee species with the habitat type (forest, grassland) and (a) bumblebee occurrence, (b) bumblebee abundance and for the interaction of the eye parameter with the forest cover in a 1 km radius of (c) bumblebee occurrence and (d) bumblebee abundance as well as the interaction of the ITD (inter-tegular distance) of bumblebee species with the forest cover in a 1 km radius of (e) bumblebee occurrence and (f) bumblebee abundance. The shaded areas represent the 95% confidence intervals.

**Figure 3:**
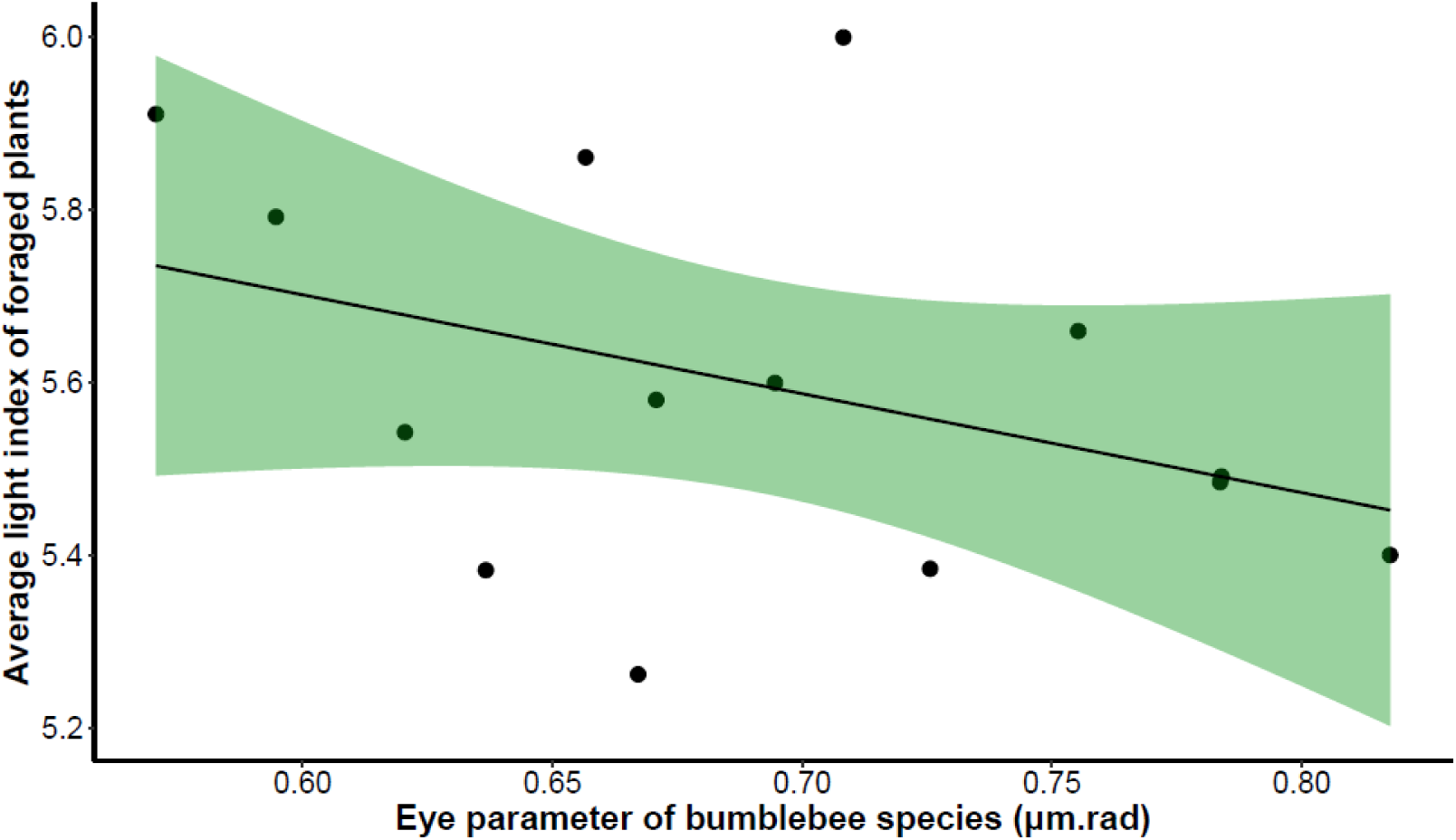
Average light index of plants foraged by bumblebee species explained by their eye parameter (workers for all species except queens for cuckoos) (Model 4). The shaded area represents the 95% confidence intervals. N.B.: The figure represents the light index of plants foraged averaged for each bumblebee species when the model accounted for the individual light index of the plant taxon.

These patterns were consistent at the community level, with the CWM of the eye parameter being higher in forest compared to grassland habitat (*Z* = -4.61, *P* < 0.001) and increasing with the forest cover (*Z* = 6.03, *P* < 0.001) (Supporting information, Figure S2).

### Bumblebees with a high light sensitivity predominantly visit shade-tolerant species

The light index of the plants that each bumblebee species forages on decreased with the eye parameter of these bumblebee species (estimate: -1.02 ± 0.36, *Z* = -2.86, *P* = 0.004). Thus, species with a higher light sensitivity are more likely to forage on plants that have a higher shade tolerance (Figure 3).

## Discussion

With this study, we found that the eyes of more common bumblebee species tended to have higher light sensitivity and rarer species tends to have lower light sensitivity. Furthermore, the species with the highest light sensitivity were more common in landscapes with increased forest cover, while the species with lowest light sensitivity were more rarely observed in landscapes with higher forest cover This pattern was also consistent at the community level with the community-weighted mean of the eye parameter being higher in forest than in grassland and increasing in both habitats together with forest cover. Moreover, bumblebee light sensitivity was related to the niche of the plants they forage on, with species with a higher light sensitivity predominantly foraging on plants with a higher shade tolerance. This strengthens our interpretation that bumblebee species both navigate and forage in habitats/landscapes that are consistent with their visual abilities, indicating that their fitness may be determined at landscape scales with consequences for their presence in local open patches. These results open new perspectives both for management and conservation measures as well as for species distribution modelling.

### Bumblebee vision and habitat use

At a coarse geographic scale, climatic conditions filter for bee species according to their temperature tolerance – estimated through body size (Hoiss et al. 2012; Oyen et al. 2016). At a landscape scale, body size, as proxy for mobility, affects where bees can forage by determining their flight range (Greenleaf et al. 2007; Zurbuchen et al. 2010; Kendall et al. 2022), and their dispersal abilities (López-Uribe et al. 2019). The eye parameter provides an insight into bumblebee habitat association while navigating and foraging. Our findings suggest that bumblebee visual abilities participate in filtering species distributions across spatial scales, by not only being involved in explaining light microhabitat use in bilberry blooming forests (Bartholomée et al. 2023), but also in how bumblebee species use their visual environment at larger scales. This is in line with results found in Sweden where bumblebee communities in grasslands had a higher light sensitivity with higher forest cover (Tichit et al. under review), and extends the range of the habitat gradient over which this trend can be observed – the pattern observed was consistent in bumblebee communities from both forests and grasslands. This increases our understanding of bumblebee species’ habitat use, as well as the robustness of predictive models, as it reduces the need to extrapolate outside of the studied habitat range, with the eye parameter being the trait responding the most to change in forest cover (Tichit et al. under review). The observed patterns could also result from eco-evolutionary dynamics (Klausmeier et al. 2020), which can happen rapidly under environmental change and be driven by plant-pollinator interactions (Pontarp et al. 2024). Bumblebee’s sensory traits and habitat use might have co-evolved such that (i) in evolutionary terms, sensory traits are affected by habitat choice and (ii) in ecological terms, sensory traits affect habitat choice and habitat-related fitness. This later point might be an important mechanism underlying bumblebee community assemblies as illustrated in this study by the landscape scale effect of forest cover. This potential co-evolution would be reinforced by the consistency between the dim light abilities of bumblebees and the shade-tolerance of the plants they forage on.

Our results were also consistent for body size – *e.g.,* larger bee species were more common in areas with higher forest cover. This can indicate both a higher mobility (Greenleaf et al. 2007; Zurbuchen et al. 2010; Kendall et al. 2022), and thus the possibility to cross larger distance of forested habitats to find more flower-rich open habitats, and a higher light sensitivity as eye size and body size are allometrically related (Jander & Jander 2002; Howland et al. 2004), and larger eyes have an overall better light sensitivity (Somanathan et al. 2007; Liporoni et al. 2020). In this perspective, future research should include a larger number of species and consider intra-specific variations in sensory traits, which might be adapted to local conditions but also allow colonies of social bees to exploit a wider range of environmental conditions. As vision was shown to influence microhabitat use in other taxa, such as butterflies (Wainwright & Montgomery 2022), upscaling these analysis to habitat and landscape levels could allow a generalisation of these patterns to other insect taxa.

### Bumblebee vision and foraged plants

Bumblebee visual abilities were related to the light niche of the plants they forage on. Species with higher light sensitivity, and thus probably a higher affinity for forest habitat and/or forested landscapes, forage on plants with a higher shade tolerance, indicating that they are able to navigate and forage in dim environments. This strengthened our understanding of the visually mediated association of bumblebee species with forest habitat and forested landscapes. This species sorting might be explained by how bumblebee visual characteristics limit their abilities to navigate through a forest to commute between habitats, but also by their ability to forage on flowers on forest understory – as species with a high eye parameter might have an advantage for both flying and foraging on dim light conditions (Spaethe et al. 2001; Baird et al. 2015). While both landscape-scale and local effects of habitat may be related to fitness differentials, the results for local scales may also represent active habitat choice by bumblebees. This study sets the basis for further investigations testing if the pattern exists for other insect taxa, for example, if the light niches of butterfly host plants is consistent with a species’ visual abilities and also if this pattern can be observed at higher trophic levels.

#### Study caveats

While accounting for the potential effects of body size, our study focused strictly on bumblebee visual abilities in relation to light intensity. By constraining the number of functional traits included in our analyses, we might have missed responses explained by other life history characteristics that likely also mediate bee’s response to environmental changes, such as diet breadth and/or nest location (Williams et al. 2010; Persson et al. 2015). We also did not include potential effects of species competition. The sample sizes for the eye parameter measures are low and from individuals captured outside the study area, which might limit the species representation (Tichit et al. under review). Moreover, we pooled the monitoring data across years, erasing inter-annual and seasonal variations. This pooled data was combined with land use data from a single year (Copernicus CLC+ Backbone 2018, (European Union’s Copernicus Land Monitoring Service information 2022)), preventing us from testing for the effects of land-use change – such as deforestation or encroachment – on the functional responses of the bumblebee species and communities. Finally, we related plant-bumblebee interactions from United Kingdom (Balfour et al. 2022) to Norwegian communities, which might not share the same plant species and we did not weight these interactions by their frequency, potentially biasing our analyses towards rarer interactions. To strengthen our analyses, bumblebee community monitoring in forest and grassland should be combined with detailed flower inventories both at the habitat and landscape scales.

### Conclusions and management perspectives

This study provides insights into the mechanisms underlying species association across scales based on their visual abilities. Bumblebees both navigate and forage in habitat and landscape according to their light sensitivity, with species and communities with higher light sensitivity being associated to dim light habitat and landscapes and foraging on plants growing in dimmer light conditions. This shows that how bees perceive and interact with their environment affects their distribution at multiple scales. We highlight the need to re-think conservation measures and management practices, most of all in forests, *e.g.* actions considering habitat light intensity together with the light sensitivity of bees and the plants they forage on. For instance, management actions to promote plants supporting threatened pollinator species in boreal forests, by favouring heterogeneous forests, with diverse stand densities and age distributions (Rhoades et al. 2018; Gómez-Martínez et al. 2020). Moreover, forest canopies provide cool microclimates (De Frenne et al. 2019; Pincebourde & Woods 2020), able to buffer heatwaves and thus being a refugia for insects, such as bumblebees which are sensitive to such extreme events (Martinet et al. 2021). Including sensory traits in species distribution modelling could allow a generalisation of the observed pattern to bee species with unknown habitat preferences but known visual trait, further improving our predictive abilities of bee responses to forest loss or the encroachment of their habitat.

## Data availability statement

The data and code will be made available on demand.

