## Supplementary material for "Forest habitat and forest dominated landscapes are associated with bumblebee species with visual traits related to light sensitivity": Table S1

**Table S1:** Bumblebee species included in the analysis of the question 1 (vision, habitat and landscape) with their total abundance in the ten years of monitoring data and the question 2 (vision and floral resources light-niche) with the number of plant taxa they were reported interacting with – and for which trait data was available. The shaded cells indicate species not included in the analysis.

| **Bumblebee species** | **Parasitic species** | **Vision, habitat and landscape** | | **Vision and floral resources light-niche** | | **Sample size**  **eye parameter measure** |
| --- | --- | --- | --- | --- | --- | --- |
|  |  | **Included** | **Total abundance** | **Included** | **Reported number of interactions** |  |
| B. bohemicus | Yes | Yes | 190 | Yes | 41 | 1 |
| B. campestris | Yes | Yes | 33 | Yes | 44 | 1 |
| B. hortorum | No | Yes | 298 | Yes | 265 | 2 |
| B. hypnorum | No | Yes | 820 | Yes | 232 | 1 |
| B. jonellus | No | Yes | 177 | Yes | 58 | 1 |
| B. lapidarius | No | Yes | 1047 | Yes | 371 | 1 |
| B. monticola | No | No | 19 | Yes | 35 | 1 |
| B. muscorum | No | No | 4 | Yes | 102 | 1 |
| B. pascuorum | No | Yes | 3797 | Yes | 455 | 4 |
| B. pratorum | No | Yes | 2457 | Yes | 314 | 1 |
| B. rupestris | Yes | No | 3 | Yes | 44 | 2 |
| B. soroeensis | No | Yes | 30 | Yes | 19 | 1 |
| B. subterraneus | No | No | 0 | Yes | 3 | 1 |
| B. sylvarum | No | Yes | 77 | Yes | 37 | 1 |
| B. terrestris* | No | Yes | 4522 | Yes | 312 | 7 |
| B. wurfelini | No | Yes | 40 | No | 0 | 1 |

#### *B. terrestris includes here both B. lucorum and B. terrestris, as both species are highly difficult to distinguish in the field (Carolan et al. 2012)**.**

Carolan JC, Murray TE, Fitzpatrick Ú, Crossley J, Schmidt H, Cederberg B, McNally L, Paxton RJ, Williams PH, Brown MJF (2012) Colour Patterns Do Not Diagnose Species: Quantitative Evaluation of a DNA Barcoded Cryptic Bumblebee Complex. PLOS ONE 7:e29251. doi: 10.1371/journal.pone.0029251
