## Supplementary material for "Forest habitat and forest dominated landscapes are associated with bumblebee species with visual traits related to light sensitivity": Table S3

**Table S3:** Post-hoc contrast analyses of the interactions between eye parameter and forest cover (%), respectively, and the habitat type (grassland, forest) for both the occurrence and abundance models (Models 1 and 2). Here we present the *emtrend* slopes together with their 95% confidence interval (CI). SD stands for standard deviation, ITD for inter-tegular distance. X indicates the absence of an interaction in the model.

| **Post-hoc contrast analysis** | **Occurrence model**  *Emtrends slopes* [95% CI] | **Abundance model** *Emtrends slopes* [95% CI] |
| --- | --- | --- |
| ***Interaction eye parameter and forest cover*** | | |
| High eye parameter (mean + SD) | 0.0067 [0.0038, 0.0096] | 0.0058 [0.0021, 0.0096] |
| Medium eye parameter (mean) | 0.0014 [-0.0008, 0.0037] | -0.0051 [-0.0078, -0.0025] |
| Low eye parameter (mean - SD) | -0.0047 [-0.0082, -0.0011] | -0.0179 [-0.0219, -0.0140] |
| ***Interaction ITD and forest cover*** | | |
| High ITD (mean + SD) | 0.0089 [0.0052, 0.0126] | 0.0075 [0.0027, 0.0123] |
| Medium ITD (mean) | 0.0010 [-0.0012, 0.0033] | -0.0060 [-0.0086, -0.0033] |
| Low ITD (mean - SD) | -0.0068 [-0.0103, -0.0033] | -0.0194 [-0.0240, -0.0148] |
| ***Interaction eye parameter and habitat type*** | | |
| Forest | 14.80 [13.60, 16.10] | 11.77 [10.33, 13.20] |
| Grassland | 10.40 [9.20, 11.50] | 7.32 [5.96, 8.68] |
| ***Interaction ITD and habitat type*** | | |
| Forest | 0.31 [0.19, 0.43] | 0.30 [0.15, 0.46] |
| Grassland | 0.14 [0.03, 0.24] | 0.08 [-0.06, 0.23] |
