## Supplementary material for "Forest habitat and forest dominated landscapes are associated with bumblebee species with visual traits related to light sensitivity": Figure S1

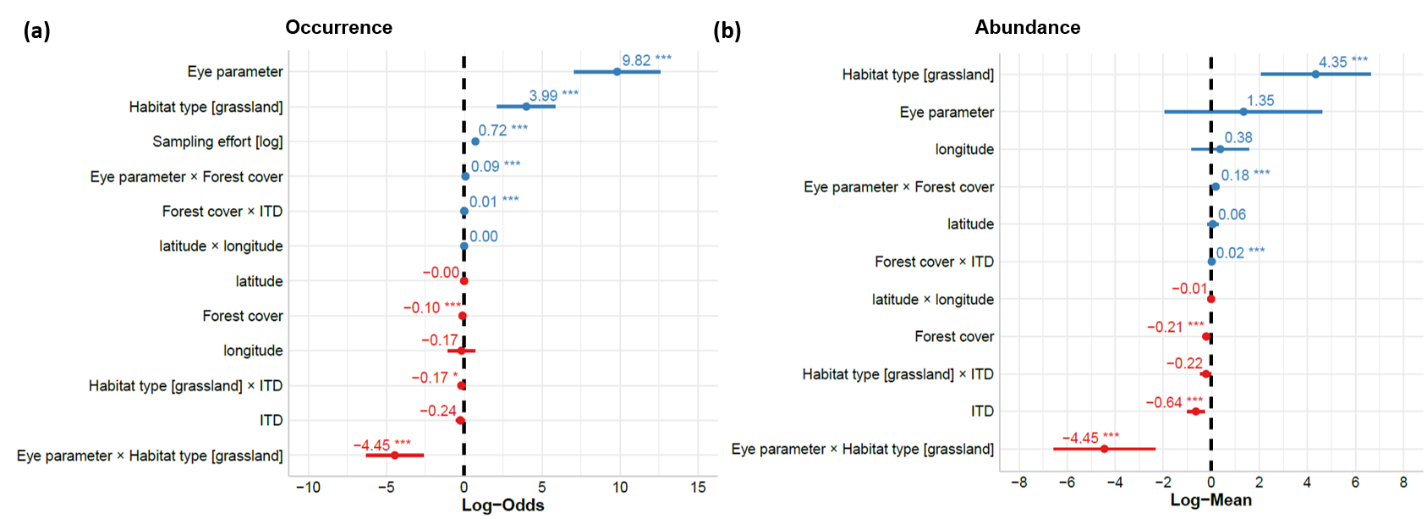

**Figure S1:** (a) Log-odd ratios of binomial Model 1 (species occurrence) and (b) log-mean generalised Poisson Model 2 (species abundance). In both models, random effects (sampling sites) were included and had a variance of 1.98*10^-4^ for Model 1 and 1.56*10^-2^ for Model 2. Blue indicates positive values, red indicates negative values. Bars indicate the 95 % confidence interval. The vertical black dashed line is the 0 value. *** indicates *p* < 0.001.
