## Supplementary material for "Forest habitat and forest dominated landscapes are associated with bumblebee species with visual traits related to light sensitivity": Figure S2

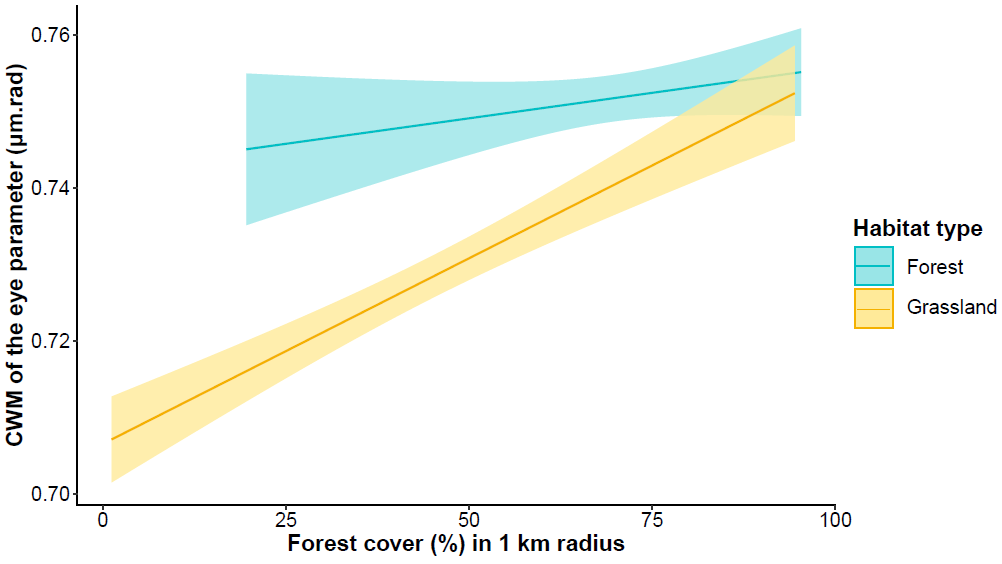

#### **Figure S2:** Community-weighted mean of bumblebee eye parameter in forests and grasslands (Model 3) (Z = -4.61, P < 0.001) along a forest cover gradient (Z = 6.03, P < 0.001) (Model 3). The shaded area represents 95% confidence interval.
